# Non-invasive characterization of oocyte deformability in micro-constrictions

**DOI:** 10.1101/2025.01.06.631295

**Authors:** Lucie Barbier, Rose Bulteau, Behnam Rezaei, Thomas panier, Elsa Labrune, Marie-Hélène Verlhac, Franck Vernerey, Clement Campillo, Marie-Emilie Terret

**Author notes:** These authors contributed equally to this work (Lucie Barbier, Rose Bulteau).

## Abstract

Oocytes naturally present mechanical defects that hinder their development after fertilization. Thus, in the context of assisted reproduction, oocyte selection based on their mechanical properties has great potential to improve the quality of the resulting embryos and the success rate of these procedures. However, the use of mechanical properties as a quantifiable selective criterion requires robust and non-destructive measurement tools. This study developed a constriction-based microfluidic device that monitors the deformation of mouse oocytes under controlled pressure. The device can distinguish mechanically aberrant oocyte groups from healthy control ones. Based on a mathematical model, we propose that deformability measurements infer both oocyte tension and elasticity, elasticity being the most discriminating factor in our geometry. Despite force transmission during oocyte deformation, no long-term damage was observed. This non-invasive characterization of mouse oocyte deformability in micro-constrictions allows for a significant advance in assessing the mechanical properties of mammalian oocytes and has potential application as a quantifiable selective criterion in medically assisted reproduction.

**Teaser:** A measurement method for translating advances in oocyte mechanics into selection criteria for improving medically assisted reproduction.

## Introduction

Mechanical parameters have been linked to the biochemical composition and structural organization of cells. Measurement of mechanical properties at the cell level can be performed by imposing a deformation within the micrometer range, using a known force applied to the cell surface. This makes the measurement of mechanical parameters a robust and non-destructive assay that reflects the cell’s physiological or pathological state (1). In assisted reproduction, oocyte quality is key for fertilization success and subsequent embryonic development (2). However, oocyte morphogenesis is error-prone, and oocytes collected during assisted reproduction are of uneven quality. Pre-selection of oocytes prior to fertilization could improve the resulting embryos quality and thus the success rate of these procedures (3). Mechanical parameters are among the promising biomarkers of oocyte quality currently under investigation (4, 5), especially as oocytes naturally present mechanical defects that hinder their development after fertilization (6).

Compared to somatic cells, oocytes are large spherical cells (80 and 120 µm in diameter for mice and humans respectively) surrounded by an extracellular matrix layer (called the Zona pellucida) only partially connected to the oocyte’s plasma membrane, creating a perivitelline space. Oocytes ready for fertilization are arrested in metaphase of second meiotic division (meiosis II) with their chromosomes aligned on the microtubule spindle. The mechanical properties of oocytes and of the Zona pellucida have been extensively studied over the last decade. The Zona pellucida has been characterized as a compressible elastic material by micro and nano-indentation experiments (7, 8). Using micropipette aspiration after removal of the Zona pellucida, it was shown that oocyte surface tension is controlled by cortical myosin-II localization and actin nucleation beneath the plasma membrane, closely related to oocyte developmental stage (9–11). Several studies have focused on assessing the subcellular mechanical properties of oocytes (12, 13), and probing of the cytoplasm with optical tweezers reveals homogeneous viscoelastic properties (14). Interestingly, the overall deformability of oocytes with their Zona pellucida under compression correlates with morphological quality criteria (15, 16), and bulk viscoelastic measurements using micropipette aspiration can be used to differentiate between viable and non-viable one-cell embryos (6, 17). In addition, the elasticity of the Zona pellucida and the viscosity of the cytoplasm are correlated with the quality of human oocytes (18–20). Furthermore, cortical tension defects affect chromosome segregation and oocyte division geometry, which have an impact on post-fertilization development (9, 11, 21). Overall, this body of work suggests that oocyte mechanical properties have a great potential as quantifiable biomarkers of oocyte quality. However, their use as a selective criterion for assisted reproduction raises the need for adapted measurement tools, and only few studies focus on a minimally invasive methodology using a platform transferable to the clinical environment (12, 15, 19, 20, 22).

Recent works have focused on the development of microfluidic-based methods enabling application of cells mechanical measurements as biomarkers for medical diagnostics. Microfluidic devices feature structures with characteristic lengths similar to those of cells, and objects can undergo controlled deformations on the micrometer scale in well-defined geometries and flows. Moreover, they are easy to use and their prototyping is versatile and inexpensive, which favors the development of new tools for medical diagnostic purposes (23). Over the past decade, microfluidic devices processing blood samples through constrictions smaller than typical cell size have been developed for the medical detection of mechanically aberrant red blood cells, as in case of malaria infection (24) or of tumor circulating cells (25, 26). In constriction-based methods, cell deformability is inferred by measuring the entry time, elongation and/or the pressure required for cell passage (26, 27). Quantitative studies have applied rheological models, such as a power law to determine single-cell elastic modulus and apparent viscosity (28–30), or Laplace’s law to determine cell surface tension (31). Constriction-based microfluidic devices could represent a promising method for assessing the mechanical properties of oocytes for medical applications. To the best of our knowledge, only two studies have proposed the measurement of oocyte deformability using micro-constrictions (32, 33). However, they both reported oocyte damage due to flow-induced shear stress during transit through the constriction (32).

In this paper, we designed a square micro-constriction with a smaller cross-section but shorter length than in previous studies. In this configuration, mouse oocytes completely fill the narrowed channel and are therefore not subjected to shear stress induced by fluid flow. We were able to measure the pressure required for oocyte passage, while assessing global and subcellular deformation. By comparing groups with known mechanical properties, we were able to distinguish mechanically aberrant oocytes from healthy control ones. Based on a mathematical model, we propose that our deformability measurements infer both oocyte cortical tension and elasticity. At the sub-cellular level, we were able to assess meiotic spindle deformation in the constriction. Importantly, after recovery, no morphological features could distinguish oocytes that passed the constriction with the off-chip control groups. Thus, we demonstrate that under reduced shear conditions, characterization of mouse oocyte deformability in micro-constrictions can be non-invasive. Our study focuses on the measurement and modeling of global deformation, and opens the possibility of a more precise analysis of the subcellular mechanical properties of oocytes in our micro-constrictions.

## Results

### Principle of the constriction-based deformability assay

In this work, we study oocyte deformation under precisely controlled pressure. To this end, we designed a constriction in a microfluidic channel that restricts the size of the oocyte in two dimensions (Fig. 1A). The restriction corresponds to a 200 μm-long segment over which the channel cross-section remains almost square but is reduced to a minimum size of 54×62 µm. Two 300 µm-long ramped segments provide a size transition between the main channels and the constriction (Fig. S1A). For deformation measurement, a single mouse oocyte is brought to the ramp segment by a moderate flow (10 µl/min). The flow is then reduced to 0 and the inlet pressure increased by 0.1 mBar every 2.5 seconds until the oocyte passes through the constriction (Fig. 1B and S1C). Once trapped in the constriction, the oocyte blocks the flow (Fig. S1D), so the applied pressure is equivalent to the inlet pressure. Moreover, oocyte elongation is only possible along the channel direction, and can therefore be measured using single-z imaging (Fig. 1B and movie S1).

**Figure 1.**
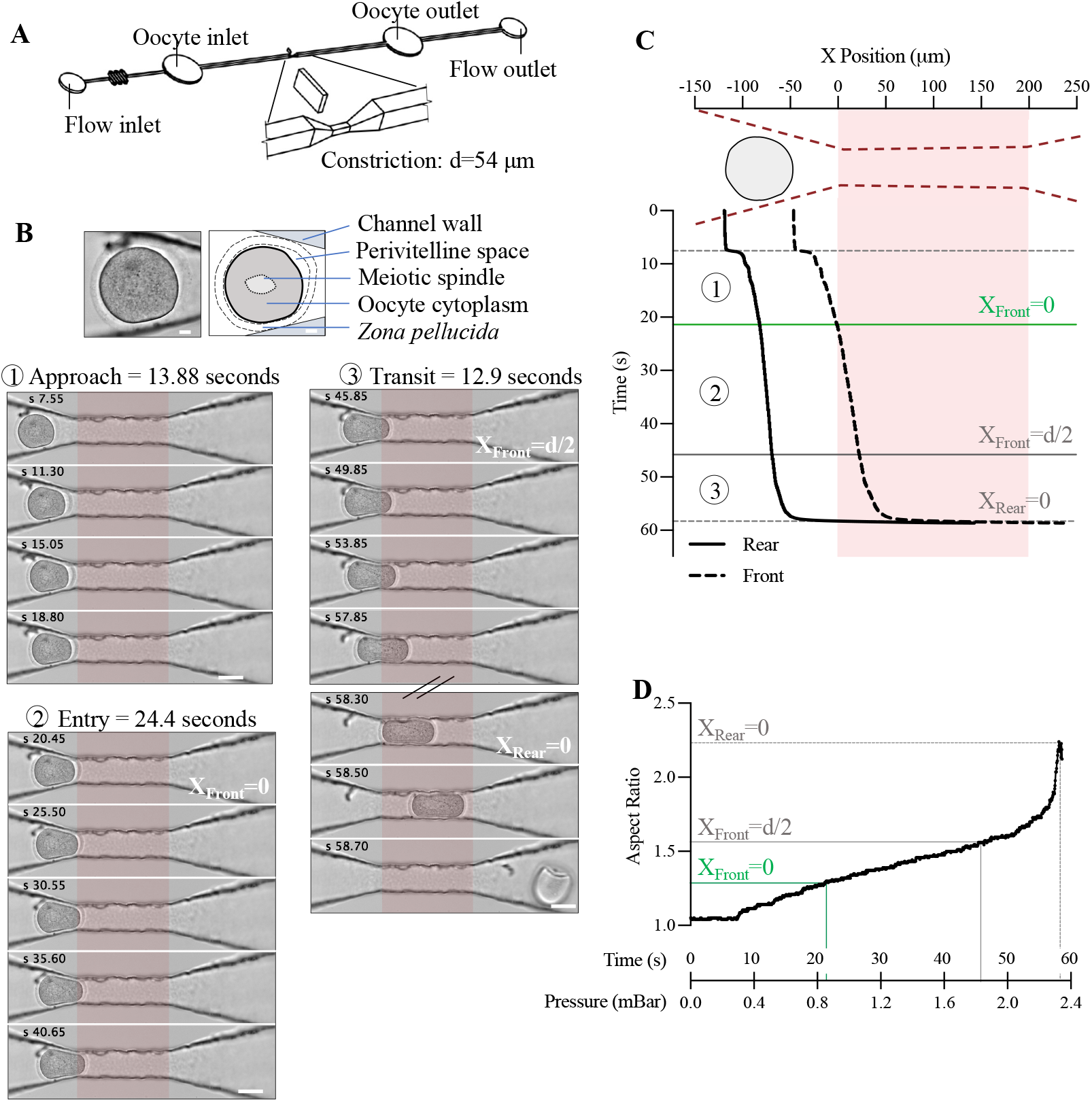
Constriction-based deformability assay for mouse oocytes. (A) 3D design of the microfluidic device. The zoom shows the 54 µm square constriction in the center of the channel. (B) Image sequence of a representative passage of an oocyte through the constriction. Upper panel: image and annotated diagram of the oocyte before entering the constriction. Scale bar = 10 µm. The bottom panel shows the three phases of deformation we have identified: approach (1), entry (2) and transit (3). The first image of the sequence corresponds to zero flow in the channel (see Fig. S1D). The images for which the front of the oocyte enters the constriction (X_Front_=0), the front of the oocyte reaches half the smallest dimension of the constriction (X_Front_=d/2), and the rear of the oocyte enters the constriction (X_Rear_=0), are annotated. Scale bar = 50 µm, time in seconds and constriction highlighted in red. (C) Front (dotted line) and rear (solid line) position of the oocyte shown in (B) relative to the start of the constriction. The drawing shows the constriction in dotted red and the initial oocyte position in grey. The constriction is highlighted in red throughout the graph. The first horizontal dotted line indicates the time of zero flow in the channel. The following lines correspond to the annotated images in (B) X_Front_=0, X_Front_=d/2 and X_Rear_=0. (D) Aspect ratio of the oocyte shown in (B) as a function of time (in seconds) since the start of the assay, and corresponding applied pressure (in mBar). The lines show the aspect ratio and pressure for the three annotated images in (B) X_Front_=0, X_Front_=d/2 and X_Rear_=0.

We used the Oocytor plugin (34) to extract the contour of the oocytes, excluding their *Zona pellucida*, from brightfield images of their passage through the constriction. We thus obtained the front and rear positions of oocytes with respect to the constriction entry (denoted by X=0 in Fig. 1C, the constriction is highlighted in red as in Fig. 1B) as well as their aspect ratio (Fig. 1D) as a function of the applied pressure. We identified three phases in oocyte deformation (Fig. 1B, C): 1) the approach phase, that starts when triggering the stepwise pressure increase and ends when the front of the oocyte enters the constriction (at which we defined the pressure P_Xf=0_). At this point, the oocytes completely block the flow under all the conditions tested. 2) the entry phase, during which the rear of the oocyte remains quasi-static while the front advances into the constriction. This phase is characterized by a linear increase in the oocyte’s aspect ratio, and we can extract the pressure at which the front of the oocyte reaches half the smallest dimension of the constriction (P_Xf=d/2_). 3) the transit phase, which corresponds to a clear displacement of the oocyte’s rear and is characterized by a rapid, non-linear increase in its aspect ratio. During this phase, we can extract the pressure at which the oocyte is completely deformed in the constriction (P_Xr=0_) (Fig. 1B, C and D).

We have thus built a tool able to monitor in-live mouse oocyte deformation as a function of applied pressure and identified three regimes associated with critical pressure values designated as P_Xf=0_, P_Xf=d/2_ and P_Xr=0_.

### Measurement of deformability identifies oocytes with aberrant mechanical properties

We hypothesized that oocyte mechanical parameters could be inferred from the measurement of pressure and aspect ratio during transit through the constriction. To test this hypothesis and identify the most discriminating parameters, we compared groups of mouse oocytes with known different mechanical properties (Fig. 2A). First, we tested healthy control oocytes at early and late stages of their first meiotic division (meiosis I), since cortical tension decreases as meiosis I progresses (Fig. 2A left diagram)(9, 10). We found no significant difference either in the pressure required for oocyte transit or in the aspect ratio inside the constriction between these two groups (Fig. 2B, C, D, Fig. S2 and movie S2). Second, we used a model of extra-soft oocytes microinjected before meiosis I, in prophase I, with cRNA encoding cortical VCA (cVCA oocytes), resulting in ectopic cortical actin polymerization via the Arp2/3 complex and myosin-II chasing from the cortex. This modification of cortical actomyosin organization induces aberrant mechanical properties, cVCA oocytes having a lower cortical tension compared to control oocytes, leading to aberrant phenotypes (Fig. 2A right diagram)(9, 11, 21). In our assay, the pressure required for cVCA oocytes to deform through the constriction (P_Xr=0_) was significantly lower than for the control oocytes, while the oocyte aspect ratio remained similar. The pressures P_Xf=0_ and P_Xf=d/2_ were also significantly lower for cVCA than for control oocytes (Fig. 2, E, F, G, Fig. S2 and movie S3).

**Figure 2.**
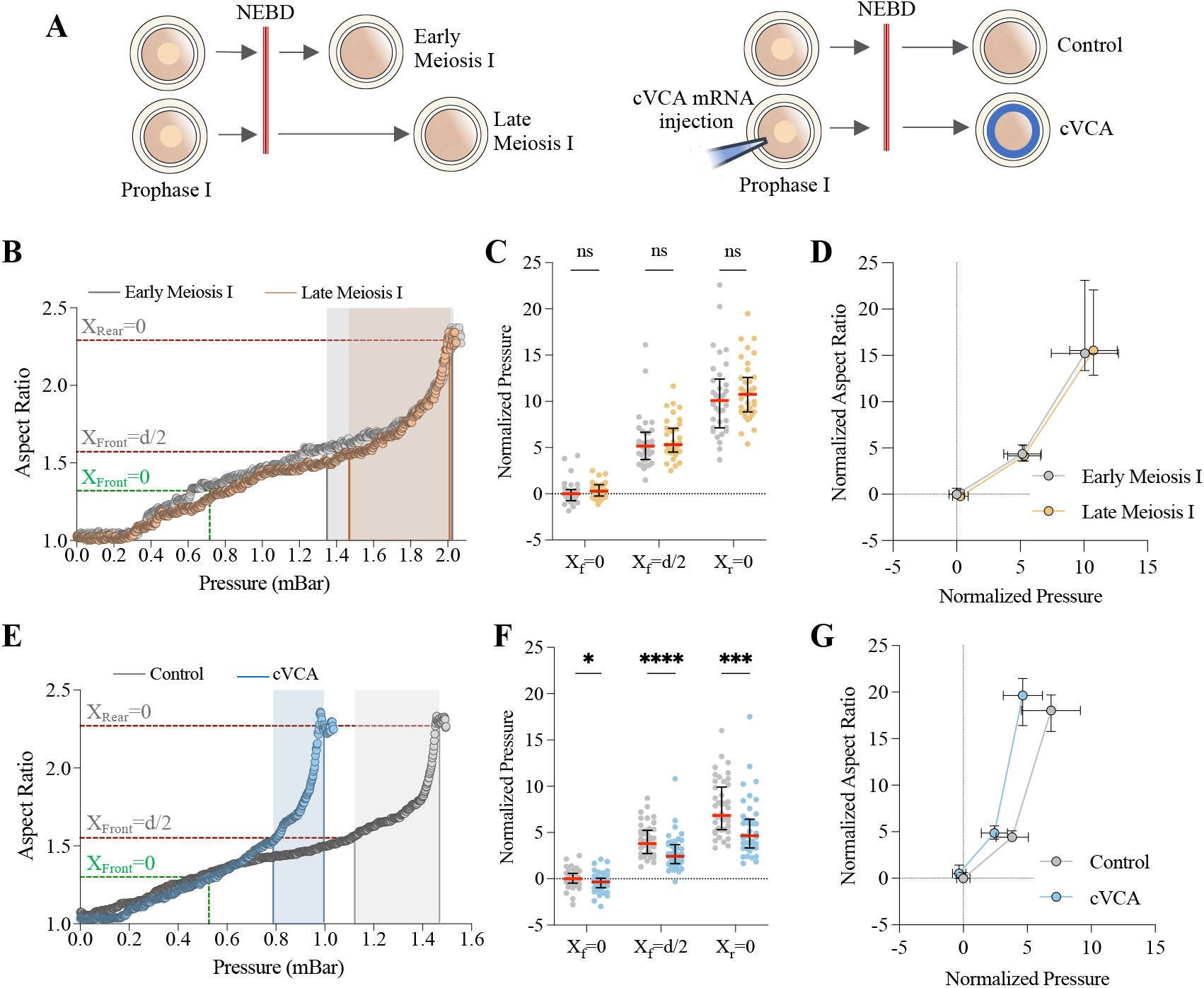
Measurement of pressure and aspect ratio during transit through the constriction for mechanically distinct groups of oocytes. (A) Diagram of methods to obtain the four oocyte groups from oocytes arrested in prophase I. The double red bar indicates the resumption of meiotic division identified by nuclear envelope breakdown (NEBD). Early meiosis I corresponds to oocytes 2 to 3 hours after NEBD; late meiosis I to oocytes 6 to 8 hours after NEBD. cVCA oocytes are obtained by microinjection of cVCA mRNA in prophase I, and measured 4 hours after NEBD. (B) Oocyte aspect ratio as a function of applied pressure for a representative early meiosis I oocyte (grey) and a representative late meiosis I oocyte (orange). The lines show the aspect ratio and pressure (in mBar) for the three critical points: X_Front_=0 (X_f_=0), X_Front_=d/2 (X_f_=d/2) and X_Rear_=0 (X_r_=0) as described in Fig. 1. The filled-in area highlights the pressure difference between X_f_=d/2 and X_r_=0. (C) Normalized pressure measured at the three critical points for early and late meiosis I oocytes and (D) Median of normalized oocyte aspect ratio as a function of the median of the applied pressure for X_f_=0, X_f_=d/2 and X_r_=0. n=34 for early meiosis I and n=35 for late meiosis I from two independent experiments. For each experiment, values are normalized to the median and interquartile range obtained for early meiosis I at X_f_=0. (E), (F) and (G): same as (B), (C) and (D) respectively for Control (grey) and cVCA oocytes (blue). n=46 for Control and n=43 for cVCA oocytes from four independent experiments. For each experiment, values are normalized to median and interquartile range obtained for control oocytes at X_f_=0. Error bars show median and interquartile range; ns: P>0.05; *P=0.0244; ***P=0.0001 and **** P<0.0001 to Mann-Whitney statistical test.

Importantly, measurements of the pressure required to pass through the constriction distinguishes oocytes with aberrant mechanical properties, but not healthy control oocytes at different stages of meiosis I. Measurement of the maximal oocyte aspect ratio was similar in all the tested conditions (Fig. S2B). For early versus late meiosis I and for cVCA versus control oocytes, differences in oocyte cortical tension have been established using micropipette aspiration (9, 10). However, other mechanical properties such as oocyte elasticity and viscosity have not been well characterized and could explain the difference in the ability to distinguish these groups of oocytes. In particular, for droplet models used to analyze micropipette aspiration experiments, surface tension governs deformability and the threshold pressure required for droplet entry in the micropipette is reached when the droplet front in the micropipette is at half the pore size (X_f_=d/2) (35). However, in our experiment, P_Xr=0_ was significantly higher than P_X*f*=d/2_ in all the conditions tested (Fig. 2D and G). This suggests that cortical tension is not the only mechanical parameter governing oocyte deformability through the constriction.

To further characterize the effect of variations in tension and elasticity on the deformability of the oocyte through the constriction, we turned to mathematical modeling.

### Tensile elastic shell modeling highlights elasticity as the determining factor in object deformation

We constructed a simplified mathematical model of the oocyte based on a classical droplet model, combined with an elastic component to represent the solid-like response of the cortex (36). The oocyte is modeled as a spherical shell with both surface tension and elastic properties. Thus, the surface stress *τ* can be written as follows:

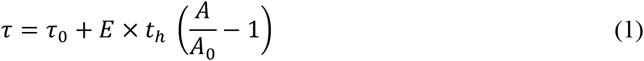

With *τ*_0_ the surface tension, *E* the elastic modulus, *t*_*h*_ the mean shell thickness, while *A* and *A*_*0*_ are the current and stress-free reference surface area of the shell, respectively. We can further idealize the oocyte shape as an axisymmetric conical geometry capped with two hemispherical shells as depicted in Fig. 3A. In these conditions, we can use Laplace’s law to obtain the following relationship between internal shell pressure *P*_*in*_ and the pressures *P*_*f*_ and *P*_*r*_ applied at the front and rear edges, respectively:

**Figure 3:**
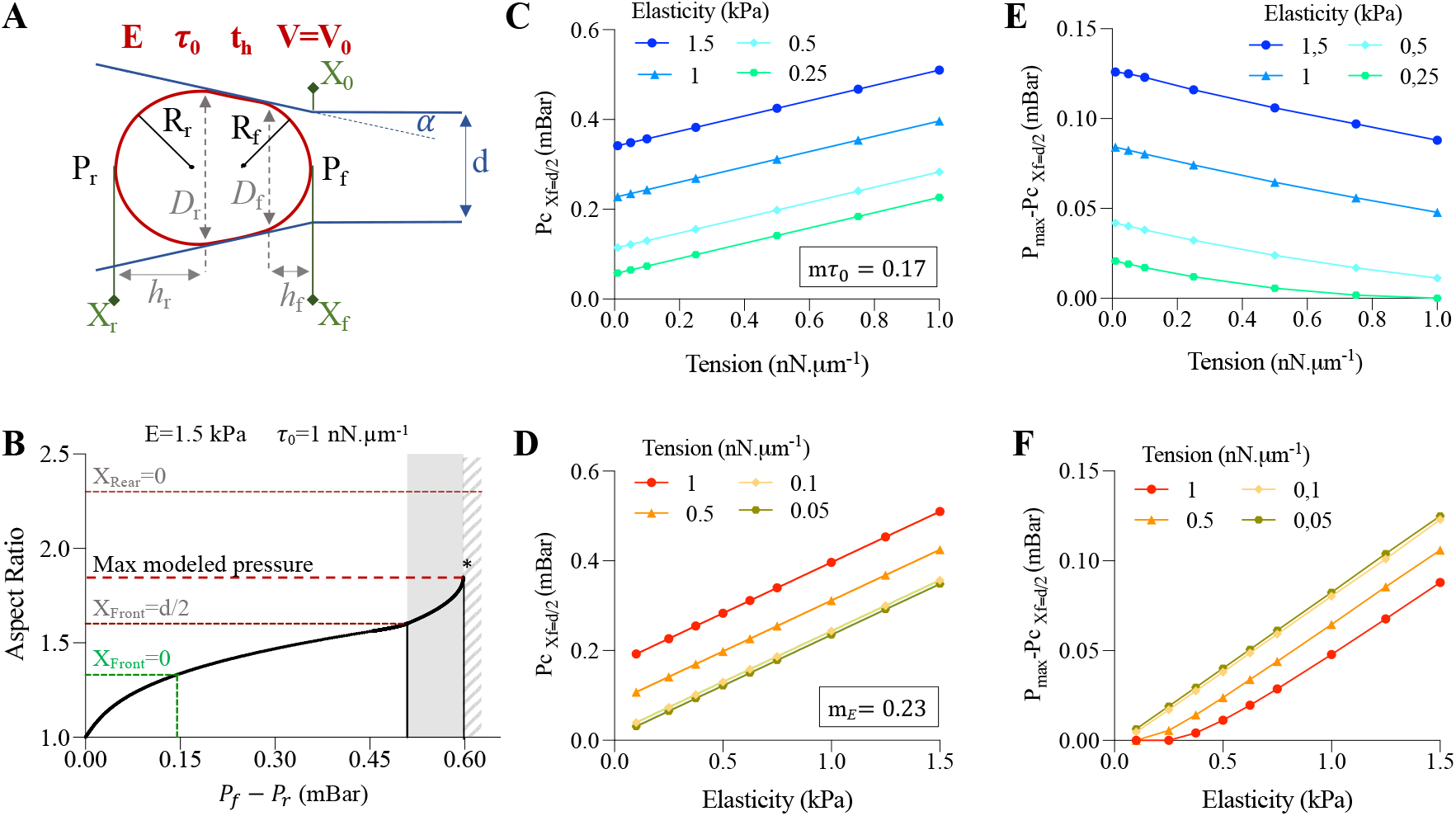
Modeling the deformation of a tensile elastic shell through the constriction. (A)Diagram of the parameters used in the mathematical model. Shell parameters are shown in red: *τ*_0_ the surface tension; *E* the elastic modulus, *t*_*h*_ the mean shell thickness and V=V0 for volume conservation. The geometric parameters of the constriction are in blue: d the minimal diameter and α the inclination of the ramped segment. Oocyte front (X_f_) and rear (X_r_) positions relative to the constriction entry (X_0_) are in green. In black: R_f_, R_r_ and P_f_, P_r_ are the radii and pressures of the front and rear edges, respectively, used to apply the Laplace law. In gray: *h*_*f*_, *h*_*r*_ and *D*_*f*_, *D*_*r*_ are the heights of the front and rear caps and the channel diameters at the base of the front and rear caps, respectively, used to calculate the equilibrium pressure difference associated with a given cell configuration (See Material and Methods). (B)Evolution of shell aspect ratio as a function of pressure gradient *P*_*f*_ − *P*_*r*_ calculated with the following parameters: D_0_=74.4 µm, E=1.5 kPa, *τ*_0_=1 nN.µm^-1^, α=9°, d=50 µm and t_h_=24.4 µm. The lines indicate the aspect ratio and pressure for the three critical points: X_Front_=0, X_Front_=d/2 and the maximum pressure P_max_ reached with the modeling. The area filled in gray indicates the pressure difference between P_Xf=d/2_ and P_max_. X_Rear_=0 corresponds to a hypothetical line and the striped area indicates the deformations observed in our experimental set-up but not reached by the model. (C)Critical pressure at X_f_=d/2 and (E) pressure difference between P_Xf=d/2_ and P_max_ as a function of shell tension calculated for different elasticity values. m*τ*_0_ indicate the slope of the linear regression for the four elasticity conditions in (C). (D)Critical pressure at X_f_=d/2 and (F) pressure difference between P_Xf=d/2_ and P_max_ as a function of shell elasticity calculated for different tension values. m_E_ indicates the slope of the linear regression calculated for the four tension conditions in (D).

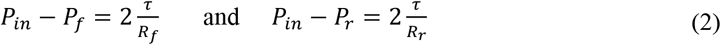

where *R*_*f*_ and *R*_*r*_ are the radii of the front and rear edges respectively (Fig. 3A). Furthermore, considering the pressure applied to the whole shell, we combine equations in (2):

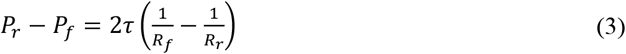

Using Eq.(1), Eq.(3) and conservation of oocyte volume (7), we can calculate the equilibrium pressure difference *P*_*f*_ − *P*_*r*_ associated with a given cell configuration (see Material and Methods). As our experimental measurement of oocyte contours excludes the *Zona Pellucida*, we corrected the model’s geometric parameters, d the constriction width, and α the inclination of the ramped segment (Fig. 3A), to obtain aspect ratio values similar to the experimental data without accounting for *Zona Pellucida* thickness (Fig. S3A). With values in the range of previous publication, E=1.5 kPa and *τ*_0_=1 nN.µm^-1^ (8–10, 12), the evolution of the shell aspect ratio as a function of the pressure difference *P*_*f*_ − *P*_*r*_ follows a similar trend to that observed in our experiments (Fig. 3B). The mathematical model shows a pressure threshold P_max_ beyond which the passage of the shell through the constriction does not require any pressure increase. For this critical pressure, the shell is not yet fully deformed inside the constriction, so the pressure P_Xr=0_ observed in our experimental set-up is not reached by the model.

If we applied Laplace’s law and considered only surface tension, the threshold pressure P_max_ would be reached when the front radius is equal to d/2 (P_Xf=d/2_). In the case of our model including an elastic component, the threshold pressure P_max_ is higher than P_Xf=d/2_ (Fig. 3B). Using this mathematical model, we could assess the effect of varying tension and/or elasticity on P_Xf=d/2_ and P_max_. We found that P_Xf=d/2_ follows a positive linear relationship with tension or elasticity, the slope being higher with variations in the shell elasticity than shell tension (Fig. 3C and D). We calculated the analytical expressions explaining the linear relationship between P_Xf=d/2_ and shell tension (slope: m*τ*_0_) or elasticity (slope: m_*E*_) as a function of our model geometry (see Material and Methods). Decreasing constriction width d increases both m*τ*_0_ and m_*E*_, while increasing ramp inclination α between 10 and 50 degrees favors m*τ*_0_ over m_*E*_ (Fig. S3B and C). Finally, the difference between P_max_ and P_Xf=d/2_ depends mainly on the elasticity of the shell and decreases only slightly with tension. As expected, P_max_-P_Xf=d/2_ tends towards 0 when elasticity decreased (Fig. 3E and F).

Overall, P_Xf=d/2_ reflects both shell tension and elasticity. In our geometry, P_Xf=d/2_ is more influenced by variations in elasticity than in tension. Moreover, the difference between P_max_ and P_Xf=d/2_ mainly reflects a variation in elasticity. A critical pressure corresponding to the modelled P_max_ is difficult to extract from our experimental measurements. However, if we approximate this threshold value by the pressure P_Xr=0_, we find that the difference between P_Xr=0_ and P_Xf=d/2_ is smaller in cVCA oocytes than in control oocytes (Fig. S3D), suggesting that cVCA oocytes have lower elasticity than control oocytes. These changes in elasticity could explain the differences in pressure measured between cVCA and control oocytes. Altogether, our tool is able to distinguish mechanically aberrant oocytes from healthy control ones, with deformability measurements inferring both oocyte tension and elasticity, elasticity being the most discriminating factor in our geometry. For medical use, our tool must be non-invasive, which we then tested.

### Meiotic spindles recover normal morphologies within hours after oocyte measurement

One of the main limitations of using constrictions to assess oocyte deformability is the shear stress-induced damage to the meiotic spindle described in previous work (32), critical for oocyte viability. To investigate the impact of our assay on the meiotic spindle, we first monitored whether it was subjected to forces during transit through the constriction in meiosis I. Spindles are visible in brightfield images as a more light-transparent area, excluding dark cytoplasmic granules. We were able to manually track the spindles during the assay, analyzing their elongation and rotation (Fig. 4A). We found that the spindles were indeed subjected to forces as they passed through the constriction, elongating by 25% along their long axis and tending to align with the channel axis (Fig. 4B, Fig. S4A). Thus, during oocyte transit through the constriction, forces are transmitted through the cytoplasm, inducing spindle deformation. Interestingly, the spindles of cVCA oocytes elongate more and are more aligned with the channel axis than those of control oocytes (Fig. 4B, Fig. S4A), suggesting an alteration in the cytoplasmic mechanical properties of cVCA oocytes, reinforcing the conclusion that our deformability assay assesses more than just differences in cortical tension. To test whether passage through the constriction induces long-term alteration of the meiotic spindle, we retrieved oocytes from the microfluidic device and assessed spindle length and chromosome alignment on the metaphase plate two hours after deformation (Fig. 4C). We found no morphological difference between meiotic spindles of control or cVCA oocytes passed through the microfluidic constriction and their unmanipulated counterparts (Fig. 4D). Finally, control and cVCA oocytes completed meiosis I, marked by polar body extrusion (PBE), at the same rate as their unmanipulated counterparts (Fig. 4E). Overall, despite transient spindle deformation during the passage through the constriction, we did not identify any long-term impact, oocytes reaching meiosis II with no visible morphological alteration.

**Figure 4:**
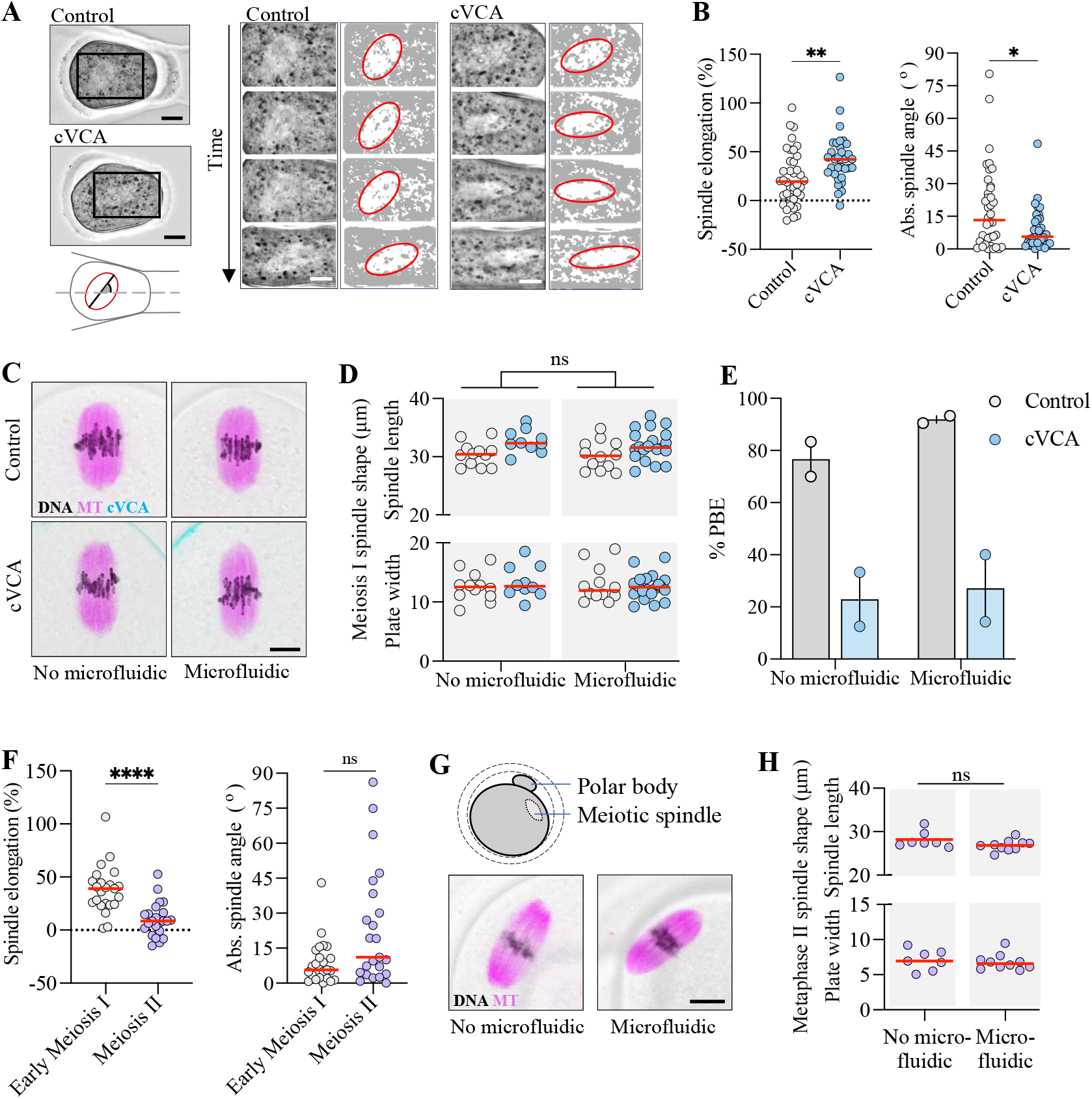
Meiotic spindle deformation and recovery after passage through constrictions. (A) Representative brightfield images of the meiotic spindles visible in a Control and a cVCA oocyte crossing a constriction. Left panel: oocyte approaching constriction; inset shows position of cropped images shown in right panel. Diagram illustrates measurements of spindle major axis and angle to channel axis, scale bars = 10 µm. The right panel shows a sequence of images cropped from the left panel. An image is shown every 15 seconds for the Control oocyte, every 10 seconds for the cVCA oocyte. The right column shows the binarized image with the spindle contour in red. Scale bars = 15 µm. (b)Left: Percentage of spindle elongation relative to its length before oocyte deformation in the constriction. Right: absolute value of the spindle long axis angle relative to the direction of the channel measured once the oocyte is fully deformed in the constriction (i.e. X_Rear_=0). n=38 for Control (grey) and n=33 for cVCA (blue) oocytes from four independent experiments. **P=0.0038 by t-test; *P=0.0388 by Kolmogorov-Smirnov statistical test. (C)Representative image of the meiotic spindle in a Control oocyte (top) and a cVCA oocyte (bottom) 2 hours after removal from the microfluidic device (left) or unmanipulated (right). Microtubule (MT) staining is shown in magenta, chromosomes (DNA) in purple, cVCA in cyan and the brightfield image is shown in light gray. Scale bar = 10 µm. (D)Quantification of spindle length measured on microtubule staining and metaphase plate width measured on chromosome staining in Control oocytes in grey and cVCA oocytes in blue. No microfluidic: n=12 for Control and n=10 for cVCA oocytes; microfluidic: n=12 for Control and n=20 for cVCA oocytes from 3 independent experiments. ns: P>0.05 for no-microfluidic versus microfluidic factor by 2-way ANOVA statistical test applied to spindle length and plate width. (E)Percentage of Control and cVCA oocytes extruding a polar body (PBE) after recovery from the microfluidic device or not manipulated. No microfluidic: n=22 for Control and n=14 for cVCA oocytes; microfluidic: n=35 for Control and n=25 for cVCA oocytes from 2 independent experiments. Each experiment is represented by a dot and the bar represent the median. (F)Same as (B) with n=23 oocytes in early meiosis I and n=23 oocytes in meiosis II from 3 independent experiments. ****P<0.0001 by t-test; ns:P=0.0591 by Kolmogorov-Smirnov statistical test. (G)Representative image of the meiotic spindle in a meiosis II oocyte 2 hours after removal from the microfluidic device (left) or unmanipulated (right). Microtubule (MT) staining is shown in magenta, chromosomes (DNA) in black and the brightfield image is shown in light grey. Scale bar = 10 µm. The diagram illustrates the organization of the oocyte in meiosis II with an extruded polar body and the meiotic spindle located below the cell cortex. (H)Quantification of spindle length measured by microtubule staining and metaphase plate width measured by chromosome staining in meiosis II oocytes: n=7 for no microfluidic and n=10 for microfluidic from one experiment. ns: P>0.05 by Mann-Whitney statistical test applied to spindle length and plate width. For all graphs, red bars represent the median.

In the standard protocols for medically assisted reproduction, hormonal treatments induce the resumption of the first meiotic division in the ovary, and oocytes that are retrieved from punctures are in meiosis II. At that stage, they are arrested in metaphase until fertilization by a sperm. Oocytes arrested in meiosis II have completed meiosis I, extruded a first polar body, and their spindle is located beneath the cell cortex, anchored in an actomyosin network. To bring our research closer to the medical application, we tested whether our microfluidic device could allow to measure and recover meiosis II oocytes without causing damage, particularly to their meiotic spindle. During transit through the constriction, the spindle of meiosis II oocytes appeared to be subjected to fewer forces than in early meiosis I. This was evidenced by the spindle being less elongated and showing a reduced tendency to align along the channel axis (Fig. 4F, S4B and Movie S5). After recovery, we found no morphological alterations in meiosis II oocytes, with no differences in spindle length nor chromosome alignment in meiosis II between those manipulated in the microfluidic devices and those that were not (Fig. 4G and H). Taken together, our results suggest that characterizing oocyte deformability in our constrictions is non-invasive, indicating potential clinical applications.

## Discussion

In this work, we describe a deformability assay using a constriction in a microfluidic device, which enables the monitoring of oocyte deformation as a function of a known applied pressure. Importantly, by measuring the pressure required to pass through the constriction, we can distinguish oocytes with aberrant mechanical properties from healthy control ones, but not healthy control oocytes at different stages of meiosis I. Based on a mathematical model combining tension and elasticity, we propose that the elastic parameter primarily governs oocyte deformability through constriction, over surface tension. At the subcellular level, we show that the meiotic spindle deforms in the constriction. However, after recovery, no morphological features distinguish oocytes that passed the constriction from off-chip control groups, suggesting that the deformability assay may be non-invasive and could be used for clinical applications.

Recent works to assess the mechanical properties of oocytes have focused on small deformations using an indenter or aspiration pipette derived from intra-cytoplasmic sperm injection protocols (17, 20). However, small deformations only probe the mechanical properties of the *Zona pellucida* and do not allow to infer the overall mechanical properties of the oocyte. On the contrary, microfluidic constrictions induce large deformations, probing the mechanical properties of the oocyte, and can easily be integrated into an on-chip platform (37). However, the only two studies that measured oocyte deformability using micro-constrictions have raised concerns about their impact on oocyte viability (32, 33). Luo *et al* designed a constriction of 50 μm wide and 150 μm high by 200 µm long and evaluated oocyte entry time under relatively high flow rates (e.g. 10 μL/min and 20 μL/min). Saffari *et al* determined the cortical tension of oocytes by measuring the pressure required for their passage through a constriction of 75 μm wide and 125 μm high by 700 µm long. Both studies were carried out on a small sample size (fewer than 30 oocytes) and reported oocyte damage: only 13 of the 23 tested oocytes reached full maturation in Saffari et al, while Luo *et al* described shear stress-induced spindle damage in 59% of tested oocytes, compared with 20% in the off-chip control groups. Our results demonstrate that constriction geometry is critical for oocyte viability. Indeed, compared to previous works, we used a constriction with a smaller cross-section but shorter length, limiting shear stress induced by fluid flow around the oocyte. As a result, we found no morphological signs of oocyte damage. Further viability tests in mice including fertilization, embryo culture up to the blastocyst stage, and reimplantation into female recipient mice, are necessary to confirm oocyte viability and developmental potential after deformation. For use in clinical applications, the assay should be further developed for use with human oocytes, and rigorously tested to ensure it is non-invasive and compatible with oocyte and embryo development.

In previous constriction-based methods, cell deformability was described either using a power-law rheological model describing viscoelastic cell behavior (28–30), or by considering cells as Newtonian liquid droplets and applying the Laplace’s law to access surface tension properties (31). Dupire *et al*. considered cells as viscoelastic droplets with surface tension generated by the actin cortex. In their model of the flowing cell, the cortical tension limits the effective stress applied to the viscoelastic material (38). In our model, we consider oocytes as incompressible spherical shells with surface tension and elastic properties, but we do not include bulk viscous properties. This minimal model was sufficient to account for oocyte deformation through constrictions. Using this modeling approach, we found that the constriction geometry is crucial to measurement sensitivity and accuracy. Indeed, the angle α of the ramp section determines the mechanical parameter that governs the oocyte entry pressure: an angle α between 10 and 50 degrees favors probing of cortical tension, while a higher or lower angle α favors probing of elasticity (Fig. S3C). Our results show that a range of mechanical parameters can be inferred in the same deformability assay with appropriate constriction geometries. This may enable to study the interplay between several mechanical parameters during oocyte morphogenesis and thus assess which parameter(s) is/are the most effective for inferring oocyte quality. In addition, we found that increasing the initial shell size (D_0_) while maintaining the same ratio of size to constriction width (D_0_/d) results in a decrease in measurement accuracy for cortical tension but not for elasticity (Fig. S3E, F). This result is essential for adapting the micro-constriction method to the measurement of larger oocytes from other species, such as human oocytes, which have a diameter of 120 µm compared to 80 µm for mouse oocytes. Thus, the shell model could be used to determine an optimal geometry that minimizes oocyte deformation while ensuring accurate measurement for oocytes selection prior to *in vitro* fertilization. To further complete these results, additional elements could be implemented in the mathematical model, including oocyte viscosity, characteristics of the *Zona pellucida*, perivitelline space, and the presence of a polar body in meiosis II oocytes, as these factors might constrain oocyte deformation and influence measurement accuracy.

Finally, one of the strengths of our approach is to combine the measurement of force and deformation with high-quality live imaging. Without specific staining, we are able to segment the subcellular elements of the oocyte as it deforms through the constriction. We present manual quantification of meiotic spindle deformation, but we were also able to observe cytoplasmic flows and thinning of the *Zona pellucida*. Further developments in image processing and analysis would be required to quantify these observations. In particular, taking into account the 3D volume of the oocyte and displacement out of the focal plane would be crucial to the accuracy of these quantifications. Nevertheless, our image data analysis adds significant information for studying load-transfer mechanisms inside the oocyte, as well as for decoupling the mechanical responses of cellular components (39). From a physical standpoint, this would allow significant advances to better describe the material properties of oocyte components. From a cell biology perspective, it would enable further studies of the cellular response induced by the mechanical load to which the oocyte may be subjected routinely in clinics during the *in vitro* fertilization procedures.

## Materials and Methods

### Mold microfabrication and scanning electron microscopy imaging

The microfluidic chip was designed on ClewinTM 5.4 (WieWeb software). The corresponding chromium mask was fabricated with the μPG 101 maskless aligner (Heidelberg Instruments Mikrotechnik GmbH) and the final silicon mold with the MJB4 alignment system (Karl Süss). To obtain 200 µm-thick channels, two 100 µm-thick photoresist dry films were successively laminated onto the silicon wafer. In a second step, the excess height of the photoresist at the constriction was removed using a CNC micro-milling machine (Minitech, Machinary Corp). To prevent the polymer sticking to the structures, the wafer was coated with a fluorinated silane by vapor deposition (trichloro 1H,1H,2H,2H-perfluorooctyl silane, Merck-Sigma-Aldrich). For scanning electron microscopy, a ten-nanometer-thick layer of gold was deposited using sputtering techniques to make the micro-milled constriction reflective to electrons. The surface morphology of the constriction was imaged by cold-field emission scanning electron microscopy at an operating voltage of 10 kV and an angle of 35 degrees.

### Microfluidic device fabrication

Liquid polydimethylsiloxane (PDMS - RTV 615, Momentive Performance) in a 1:10 base/crosslinker ratio was poured onto the custom mold, degassed and cured for at least 2 hours at 70°C. After cutting the PDMS pieces and punching the inlets and outlets with a biopsy punch (Electron Microscopy Sciences, Hatfield), the PDMS pieces were cleaned by sonication in isopropanol for 30 seconds and left to dry overnight. The PDMS pieces were then bonded to a microscopy glass slide after O2 plasma treatment of both surfaces (50 W for 1 minute, O2 flow rate 20 sccm, pressure 0.15 torr, Cute plasma oven, Femto Science), and left for 5 minutes at 90°C to ensure bonding. The microfluidic devices produced were connected to a pressure controller and flow sensor (Fluidgent), filled with culture medium and left for at least 1 hour to equilibrate at 37°C before measuring oocytes.

### Mouse oocytes collection and culture

Oocytes were extracted by shredding ovaries collected from 8 to 11-weeks-old OF1 female mice (Charles River Laboratories) in homemade M2 medium (40) supplemented with 1 μM milrinone (41) to synchronize them in prophase I. Meiosis resumption is triggered by transferring oocytes into milrinone-free homemade M2 medium.

All live culture and imaging were carried at 37 °C and under oil apart for microfluidic measurement.

All animal studies were performed in accordance with the guidelines of the European Community and were approved by the French Ministry of Agriculture (authorization D750512).

### Plasmids, *in-vitro* transcription of cRNAs and oocyte microinjection

As previously described (11), we used the following construct pRN3-EzTD-mCherry-VCA. *In vitro* synthesis of cRNAs was performed using mMessage mMachine kit (Ambion) and subsequent purification with the RNeasy kit (Qiagen) following the manufacturer’s instructions. cRNAs were centrifuged at 4 °C for 45 min at 20,000 g before microinjection. cRNAs were injected in prophase I arrested oocytes using an Eppendorf Femtojet micro-injector. cRNA translation was allowed for 2h before meiosis resumption was triggered.

### Oocyte measurement in the constriction

After deposition in the inlet, the oocyte was driven to the constriction at a moderate flow rate of around 10 μl/min. Once the oocyte reached the constriction, the flow rate was adjusted to 0 μl/min to stop the oocyte in contact with the channel walls without being deformed. The inlet pressure was increased by 0.1 mBar every 2.5 seconds until the oocyte passed through the constriction. Pressure and flow were recorded every 50 ms for the duration of the oocyte’s passage. Automation and recording were performed using OxyGEN software (Fluidgent). An image of the oocyte was taken every 50 ms on an inverted brightfield microscope (Diaphot 300, Nikon) with a 20X dry objective and a CMOS camera (ORCA-Spark, Hamamatsu). Triggering of image recording and stepwise pressure increase were synchronized using auto-click software (Fig. S1B). The contour of the oocytes, excluding the *Zona pellucida*, were obtained using Oocytor plugin (34). The contours were then analyzed using Fiji software (NIH) to extract oocyte position and aspect ratio as a function of time since the triggering of stepwise pressure increase and image recording. Due to the experimental variation observed in the P_Xf=0_ measurement between control conditions (Fig S2A), all pressure values were normalized to the median and interquartile range of the P_Xf=0_ measurement for the control condition of each experiment.

### Determination of the equilibrium pressure for a given cell configuration

#### Cell geometry

To find the equilibrium pressure difference Δ*P* associated with a given cell configuration, we assume that the cell geometry is composed of three different sections (Fig. 3A): (a) a spherical cap on the rear end with a volume 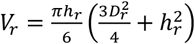, where *h*_*r*_ is the height of cap and *D*_*r*_ is the diameter of the channel at the base of the cap, (b) a partial cone with a volume 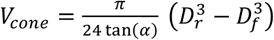 where *D*_*f*_ is the diameter of the channel at smaller base of the partial cone, and (c) a spherical cap on the front end with a volume 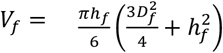 with *h*_*f*_ the height of the cap. The volume of the cell is then calculated as:

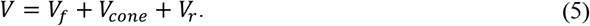

These volumes can be computed from the channel geometry and the volume conservation condition that require *V* = *V*_0_ (where *V*_0_ is the initial volume of the cell). To finally account for the cortex elasticity (in equation (1)), we evaluate the total surface area of the cell, composed of two spherical caps and a partial cone, i.e.:

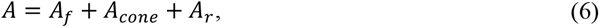

where

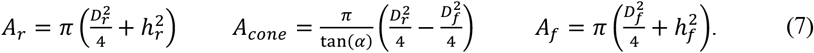

It should be noted that these formulations are valid before the front spherical cap becomes a hemisphere (*P*_*Xf*=*d*/2_). After this stage, the shell will be composed of four geometrical sections, including two spherical caps, a partial cone and a cylinder. We can show however, that at that stage the motion of the cell is unstable (i.e. cell translation is associated with a drop in pressure difference). Since results presented in this work only relate to the stable branch of the pressure/motion curve, this stage is not considered in our analysis. In the analysis, we finally assume that the cell is initially a perfect sphere, the initial volume and surface area of the shell are thus 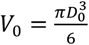 and 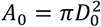, where *D*_0_ is the cell’s initial diameter.

#### Numerical procedure

The solution to the problem relies on finding the relationships between the height and radius of the bases of the caps. Thus, the solution procedure is implemented into two successive stages. a) In the first stage, the wall of the tapered section of the microchannel is identical to the tangent lines to both spherical caps at the point of contact. With this assumption, the heights of the caps are related to the radii of the bases as follows:

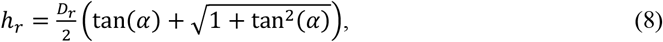

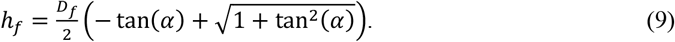

This stage lasts until the wall of the tapered section of the microchannel is no longer tangent to the downstream spherical cap. This corresponds to *X*_*f cap*_ = *X*_*f*_ − *h*_*f*_ = *X*_0_. b) In the second stage, the downstream cap enters the constriction part of microchannel (*D*_*R*_ = *d*). Thus, *h*_*R*_ is independent from *D*_*R*_ and Eq. (9) is no longer valid. However, the tangency condition still holds for the upstream cap, and we again use Eq. (8). Note that in contrast to the experimental procedure, the numerical procedure consists of finding the pressure difference associated with a prescribed cell location therefore avoiding numerical convergence issues related to the problem’s inherent loss of stability.

The numerical procedures for the two aforementioned stages are as follows:

#### First stage

1. For a given value of *X*_*f cap*_, we calculate *D*_*f*_ = −2 tan(*α*) (*X*_*f cap*_ − *X*_0_) + *d*
2. We find *h*_*f*_ by Eq. (9)
3. Given that *D*_*r*_ = −2 tan*(α*) (*X*_*r cap*_ − *X*_0_) + *d* and Eq. (8), we find *X*_*r cap*_ that satisfies the incompressibility condition
4. We calculate the radii of the spherical caps as 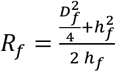 and 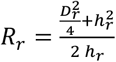
5. We find the surface area *A* and surface force *τ* are calculated by Eq. (6) and Eq. (1), respectively
6. Aspect ratio is calculated by 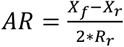
7. We calculate *P*_*r*_ − *P*_*f*_ by equation (4)
8. We repeat steps (1) through (7) until *X*_*fcap*_ = *X*_0_

#### Second stage

1. We set *D*_*f*_ = *d*
2. For a given value of *h*_*f*_, we solve the incompressibility equation to find *X*_*r cap*_. To solve this equation, we again use Eq. (8) and *D*_*r*_ = −2 tan*rα*) *rX*_*r*_ − *X*_0_) + *d*)
3. We calculate the radii of the spherical caps as 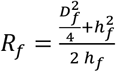and 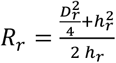
4. We find the surface area *A* and surface force *τ* are calculated by Eq. (6) and Eq. (1), respectively
5. Aspect ratio is calculated by 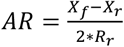
6. We calculate *P*_*r*_ − *P*_*f*_ by equation (4)
7. We repeat steps (2) through (6) until *h*_*f*_ = *d*/2

The numerical procedure were done on MATLAB software (MathWorks), the code is available on https://github.com/Behnam-Rz/Oocyte_Constriction/tree/main

### Analytical closed form for the critical pressure

Given the relationships between geometry parameters, we can find an analytical closed form for the critical pressure *P*_0_ = *P*_*Xf*=*d*/2_. This pressure corresponds to the last iteration of the second stage simulation procedure. By setting 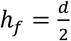 and *D*_*f*_ = *d*, and accounting for the incompressibility equation, the diameter 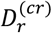 of the base of the upstream cap is calculated as:

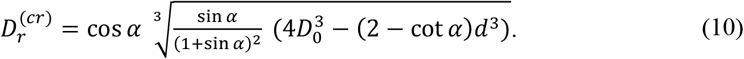

Therefore, the surface area of the shell at *P*_*Xf*=*d*/2_ is calculated as follows:

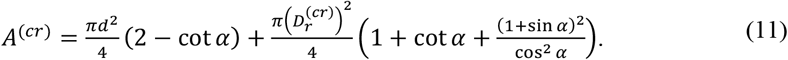

At this pressure, the radius of the downstream cap is equal to the constriction radius 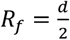 and the radius of the upstream cap (using trigonometric identities) is calculated as 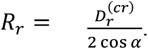. Combining these formulas, the close form formula for *P*_*Xf*=*d*/2_ is calculated as:

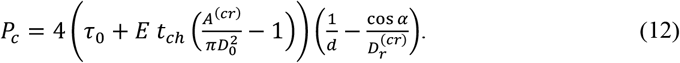

The above equation can be rewritten as:

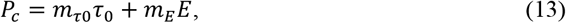

where 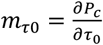shows the sensitivity of the critical pressure to the surface tension and 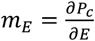 measures the sensitivity of the critical pressure to the elasticity of the shell. Comparing Eq. (12) with Eq. (13), we find:

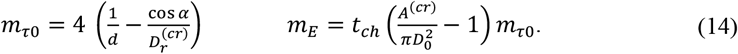

Thus, Eq. (13) shows that the critical pressure changes linearly with both surface tension and elasticity (Fig. 3C, D). Furthermore, Eq. (14) can be used to study the effects of determining parameters, such as the constriction diameter *d* (Fig. S3B), the ratio *D*_0_/d (Fig. S3E, F), and the inclination angle *α* (Fig. S3C) on *P*_*Xf*=*d*/2_.

### Meiotic spindle length and chromosomes alignment

Oocytes were incubated 30 minutes with 0.1 μM SiR-tubulin (SiR-tubulin Kit, Spirochrome) and Hoechst 5 ng/mL (Sigma-Aldrich H6024) to label microtubules of the meiotic spindle and DNA of the chromosomes. Images were acquired on a confocal-spinning disk microscope (DMI6000B, Leica and CSU-X1 Spinning disk, Yokogawa) enclosed in a thermostatic chamber (Life Imaging Service) using a Plan-APO 40x/1.25 NA objective and a CCD camera (Retiga 3, QImaging). Metaphase plate width and meiotic spindle length were measured by manually placing bounding boxes on DNA fluorescent signal and microtubule fluorescent signal respectively using the Fiji software (NIH). Measurements were done only on spindles parallel to the imaging plane (21).

### Statistical Analysis

Appropriate statistical test was applied according to data normality (determined by Shapiro-Wilk and Kolmogorov-Smirnov tests) and content (paired observations and/or equal variance). Respective tests and *n* numbers are indicated in the figure legends. Statistical analysis was performed using GraphPad software (PRISM).

## Acknowledgments

We thank Elvira Nikalayevich and Maria Almonacid for feed-back on the manuscript; Anastasia Shihabi for cRNA production and all the members of the Verlhac-Terret laboratory and the LAMBE for support in developing the project. The microfluidic devices fabrication was performed at the IPGG technological platform supported by the CNRS (UAR 3750).

## Author contributions

CC and MET conceived the project that was directed by CC, MET and LB. LB and RB designed, performed and analyzed all experiments with the help of EL. BR and FV built the mathematical model. MHV was involved in some aspects of project supervision. TP participated in the engineering of the chip. LB wrote the manuscript, which was seen and corrected by all authors.

## Data and materials availability

All data are available in the main text or the supplementary materials (optical image sequences, flow and pressure recording) are available upon request to the corresponding authors. Data are archived at Collège de France. The Matlab code used for determination of the equilibrium pressure for a given cell configuration is available from: https://github.com/Behnam-Rz/Oocyte_Constriction/tree/main

