## Supplementary materials for "Non-invasive characterization of oocyte deformability in micro-constrictions"

Lucie Barbier *et al.*

**This PDF file includes:**

Figs. S1 to S4
Movies S1 to S5

**Other Supplementary Materials for this manuscript include the following:**

Movies S1 to S5

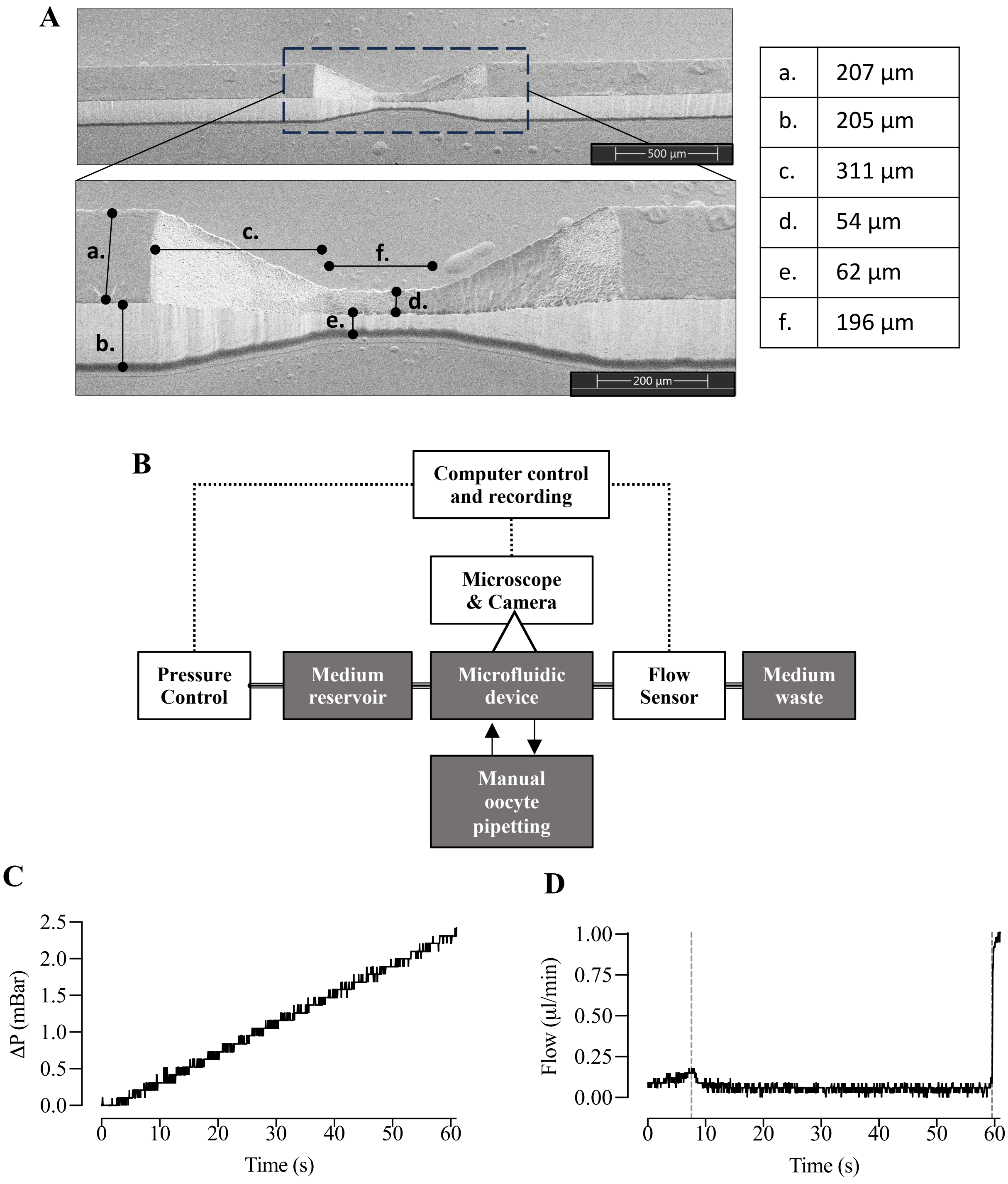


| | |
|---|---|
| a. | 207 µm |
| b. | 205 µm |
| c. | 311 µm |
| d. | 54 µm |
| e. | 62 µm |
| f. | 196 µm |

**Fig. S1.**

(A) Electron microscopy image of the silicon wafer used to mold the microfluidic device.

(B) Diagram of the components used for the deformation assay. The pressure controller, camera and flow sensors are connected to the computer for process control and recording.

(C) Pressure applied to the inlet of the microfluidic device for oocyte passage illustrated in Fig. 1B.
(D) Flow rate measured at the outlet of the microfluidic device for oocyte passage illustrated in Fig. 1B. Dotted vertical lines indicate when the oocyte inserted into the constriction stops the flow and when it is released.

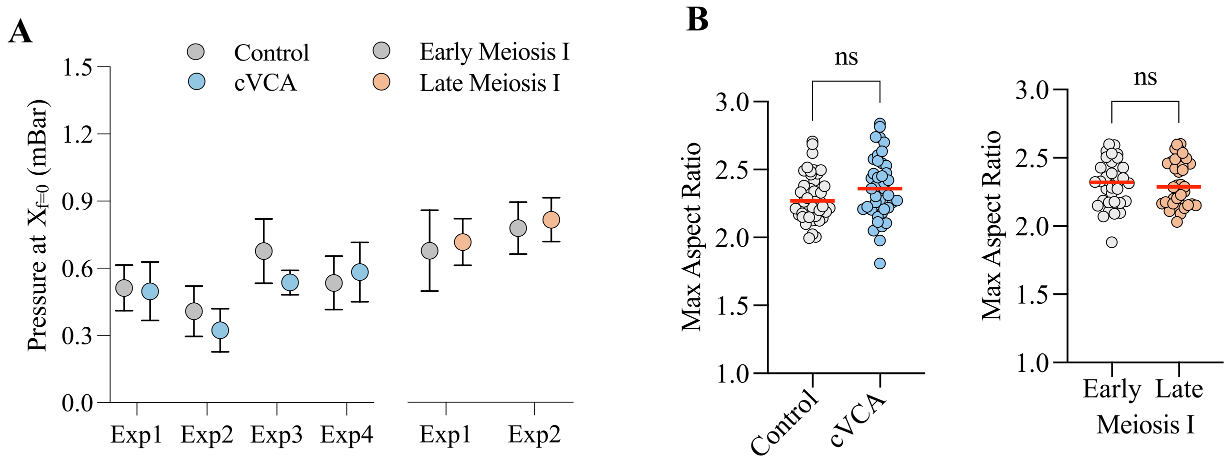


**Fig. S2**

(A) Pressure measure at $X_{f}$=0 for each independent experiments. Median and interquartile range are shown.

(B) Maximum aspect ratio measured for Control, cVCA, early and late meiosis I oocytes. ns: P>0.05 to Mann-Whitney statistical test.

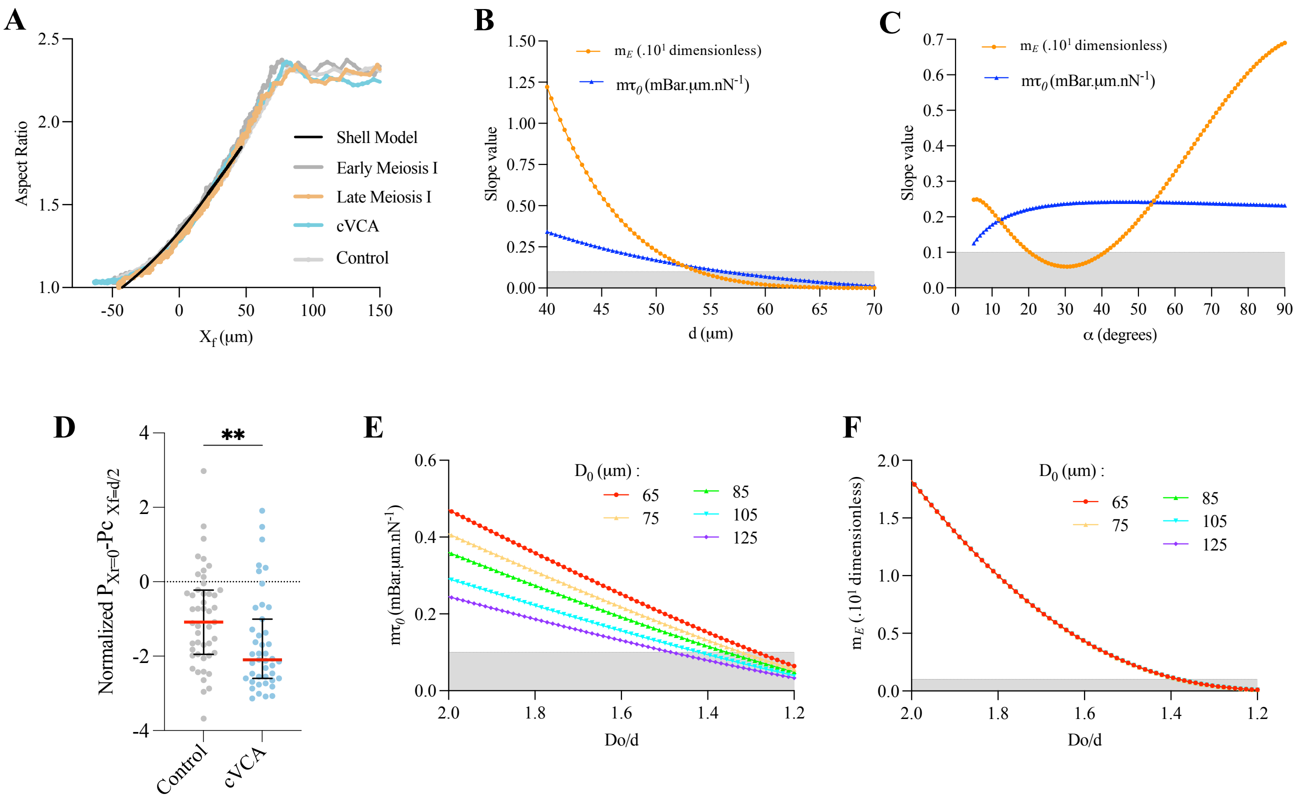


**Fig. S3.**

(A) Aspect ratio of oocytes in Fig. 2B and E as a function of the position of the oocyte front relative to the constriction entry. The black line shows the modeled values obtained with the following parameters: $D_0$=74.4 µm α=9°, d=50 µm.

(B) (C) Value of the slopes $m\tau_0$ and $m_E$ as a function of constriction diameter d (B) or ramp section angle α (C) calculated for $D_0$=74.4 µm, E=1.5 kPa, $\tau_0$=1 nN.µm$^{-1}$ and $t_h$=$(D_0 - d)$.

(D) Difference between normalized pressure measured at $X_f$=d/2 and $X_r$=0 and shown in Fig. 2F. Error bars show median and interquartile range; **P=0.0033 to Mann-Whitney statistical test.

(E) (F) Value of the slopes $m\tau_0$ (E) and $m_E$ (F) as a function of the ratio between initial shell size and constriction diameter ($D_0$/d) for different values of $D_0$. In (F) the curves overlap for all tested values of $D_0$.

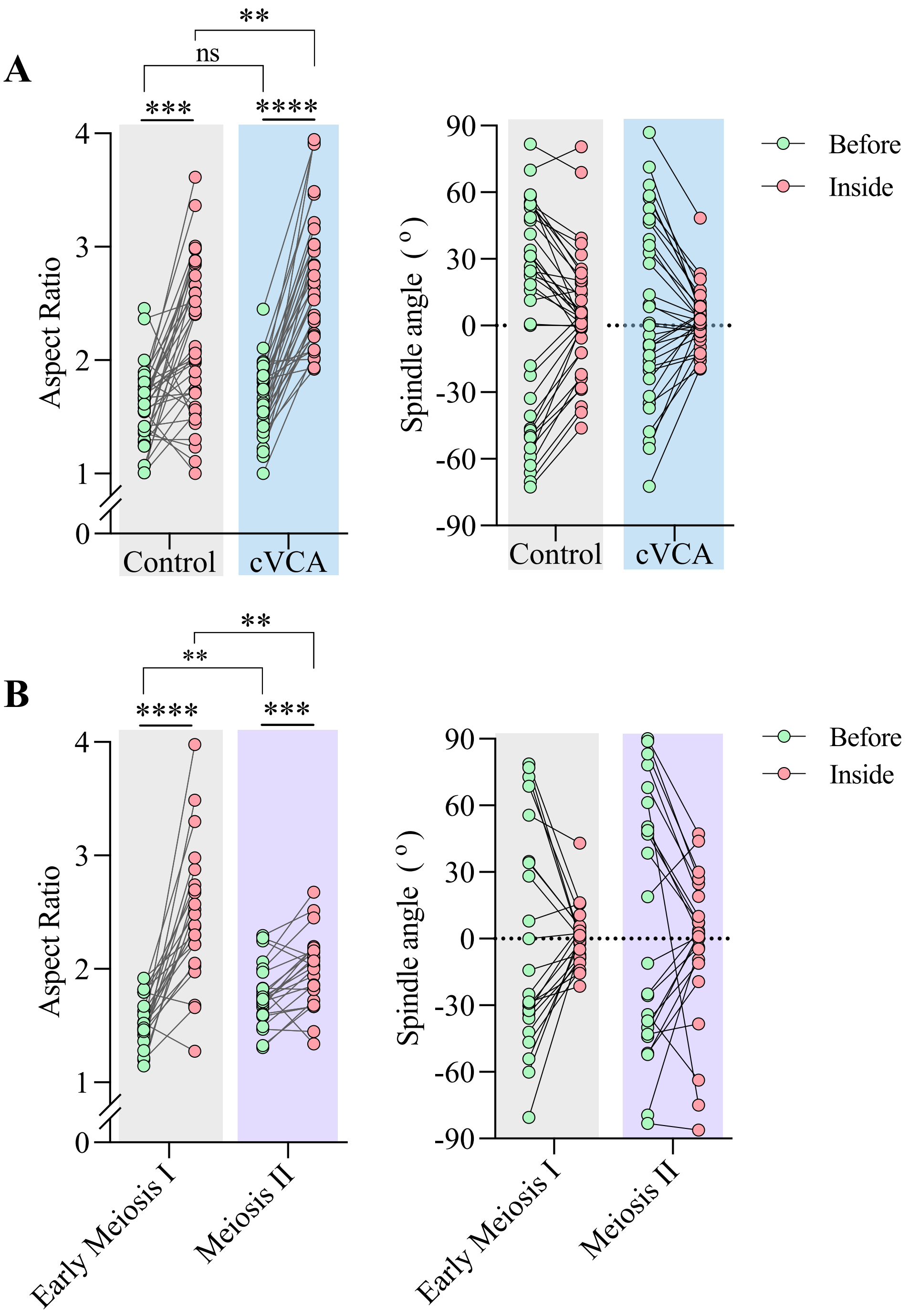


**Fig. S4.**

(A) Meiotic spindle aspect ratio (left) and angle of spindle major axis to canal direction (right) before constriction and once the oocyte is fully deformed inside the constriction (i.e. $X_{Rear}$=0). n=38 for Control oocytes and n=33 for cVCA oocytes from four independent experiments.

(B) Same as (A), n=23 for oocytes in early meiosis I and n=23 for oocytes in meiosis II from 3 independent experiments.

ns: $P>0.05$ and **$P<0.01$ by t-test with Welch correction in (B); ****$P<0.0001$ and ***$P<0.001$ by paired t-test.

**Movie S1 (separate file).**
Passage of the oocyte through the constriction as shown in Fig. 1B. The movie is cropped to start when the flow is zero in the channel. The three phases of deformation are highlighted in red for approach, green for entry and magenta for passage. Time in seconds, scale bar = 40 µm.

**Movie S2 (separate file).**
Passage of a representative early meiosis I oocyte (black; top) and late meiosis I oocyte (orange; bottom) through the constriction. Same cells as in Fig. 2B. Time in seconds, scale bar = 40 µm.

**Movie S3 (separate file).**
Passage of a representative Control oocyte (grey; top) and cVCA oocyte (blue; bottom) through the constriction. Same cells as in Fig. 2E. Time in seconds, scale bar = 40 µm.

**Movie S4 (separate file).**
Passage of a representative Control oocyte (black; left) and a cVCA oocyte (blue; right) through the constriction. Same cells as in Fig. 4A. Time in seconds. The walls of the channel were erased using mean gray value filters and the movie was adjusted according to the rear position of the oocyte to better visualize the position and shape of the meiotic spindle. The measurements shown in Fig. 4B were performed on unmodified movies. Scale bar = 20 µm.

**Movie S5 (separate file).**
Passage of a representative early meiosis I oocyte (black; left) and a meiosis II oocyte (magenta; right) through the constriction. Time in seconds. The walls of the channel were erased using mean gray value filters and the movie was adjusted according to the rear position of the oocyte to better visualize the position and shape of the meiotic spindle. The measurements shown in Fig. 4F were performed on unmodified movies. Scale bar = 20 µm.

For all movies, one image is shown every two acquisitions.
